# Interpretation of murine models of cardiac disease is enhanced by consideration of divergently expressed human-mouse orthologous genes

**DOI:** 10.1101/2024.06.15.599153

**Authors:** Oisín Cappa, Chris Watson, David Simpson

## Abstract

Humans and mice share remarkably similar cardiac anatomy and physiology and have many orthologous genes with high sequence similarity. Together with their small size and short lifespan, these factors are major contributors to the popularity of mice as research models. Nonetheless, findings in mice often fail to translate into humans. We propose that this may be due in part to differences in gene expression. Advances in molecular genomics have enabled profiling of gene expression at the level of individual cells through single-cell RNA-sequencing. Large repositories of human and murine data across many organs and tissues, including the heart, are now available. However, there is currently a lack of comparative research which directly quantifies similarities and differences in expression of orthologous genes between human and mouse.

This study is the first, to our knowledge, to directly compare celltype-specific expression conservation and divergence between orthologous genes in human and mouse heart. Cardiac data from 7 human and mouse scRNA-seq publications was retrieved, with mouse data converted to equivalent human gene orthologues. The datasets were integrated and subjected to comparative transcriptomic analysis in R using Seurat.

Classical celltype markers did not perform equally across species. We have identified species-conserved celltype marker genes, providing a useful resource to identify cardiac cell populations in both species. Further, we have identified and ranked divergence between species for each cardiac celltype, highlighting genes which vary considerably in expression between human and mouse heart. Many of these have been genetically modified to create mouse models or encode proteins targeted by pharmaceutical drugs. Researchers should carefully consider these highly divergent genes when planning, or interpreting data from, mouse models of human biology.

## Introduction

Humans and mice share remarkably similar cardiac anatomy and physiology. Coupled with practical advantages such as a short reproductive cycle, rapid development, and the availability of genetically modified strains, this makes mice an excellent model for studying cardiovascular disease, resulting in thousands of publications annually recorded on PubMed and Web of Science. Humans and mice also share a very similar genetic background; approximately 90% of both genomes exist within syntenic blocks (Mouse Genome Sequencing Consortium, 2002) and between 70 and 80% of protein-coding genes have one-to-one gene orthology, with approximately 85% sequence similarity on average (Breschi, Gingeras and Guigó, 2017). However, it is becoming increasingly apparent that findings in these mouse models often do not translate to the human situation. We hypothesise that this may be attributed to differences in expression of the orthologous genes within the human and mouse tissue.

High-throughput RNA-Sequencing techniques that measure genome-wide gene expression make it possible to compare mouse and human tissues at the molecular level to assess conserved and divergent aspects of gene expression (Zheng-Bradley *et al,* 2010, Cardoso-Moreira *et al*, 2019). Furthermore, the development of single-cell RNA sequencing (scRNA-seq) has revolutionised the field of transcriptomics by enabling the measurement of gene expression within individual cells (Macosko *et al*, 2015), and therefore the comparison of orthologous gene expression at the celltype level. The exponential increase in studies profiling all cellular populations within different organs has led to the development of comprehensive cell atlases across several species, primarily human and mouse, which serve as valuable searchable references of celltype-level gene expression. Notable groups managing human single-cell expression atlases include Tabula Sapiens (The Tabula Sapiens Consortium, 2022) and Human Cell Atlas (Regev et al, 2017) consortia, whichhave assembled data for over 30 organs and tissues, across more than 200 publications. Resources of comparable aim and scope also exist for mice such as Mouse Cell Atlas (Wang *et al*, 2022) and Tabula Muris (The Tabula Muris Consortium, 2018).

The well-defined gene orthology between human and mouse makes it possible to directly compare gene expression between species by restricting the analysis to orthologous genes. A commonly used approach to compare cross-species gene expression involves searchable databases of gene expression and/or celltype marker genes, calculated independently within datasets. Large platforms for such information include the UCSC Cell Browser (Speir *et al*, 2021) and EMBL-EBI Expression Atlas (Papatheodorou *et al,* 2020). By searching data for tissues across multiple species, broad similarities or differences can be discerned for individual genes of interest. However, this approach is limited by low throughput and the absence of *quantitative* cross-species comparisons to provide meaningful insights into relative levels of expression.

This field is developing rapidly, such that recent works now recognise the importance of directly integrated cross-species datasets. This approach has been used in the cross-species investigation of neocortex development (Li et al, 2020), cellular composition changes in Alzheimer’s disease (Johnson *et al*, 2020), and immune cell heterogeneity in atherosclerosis (Zernecke *et al*, 2023), among others. Of particular relevance is the recent publication on cross-species transcriptomic dysregulation in heart failure (Jurado *et al,* 2024), which identifies common and divergent regulatory pathways implicated in this cardiac disease. However, to our knowledge, no celltype-level resource has yet been published for the comparative assessment of the human-mouse transcriptome in healthy heart.

In this study, we present an approach to directly integrate healthy human and mouse scRNA-seq cardiac datasets. By focusing on orthologous genes contrasted within equivalent celltypes, we characterise interspecies conservation and divergence within the human and mouse heart. Our aim is for this resource to improve the planning and interpretation of cross-species transcriptomic investigations.

## Methods

### Dataset Retrieval and Human-Mouse Orthologue Conversion

Publicly available human and mouse cardiac single-cell RNA sequencing datasets were retrieved from genomic data repositories, restricted to healthy adult single-cell/single-nuclei data derived from 10x Genomics (Pleasanton, CA, USA) (droplet-based) gene expression profiling. Murine data from five publications were accessed, from Skelly *et al*, 2020 (E-MTAB-6173), McLellan *et al* 2020 (E-MTAB-8810), Galow *et al*, 2020 (E-MTAB-8751, E-MTAB-8848), Wu *et al*, 2021 (GSE180821) and The Tabula Muris Consortium, 2018 (GSE132042). In total, data from 17 adult mouse hearts were processed, including 56,856 single cells. Human data was accessed from the Heart Cell Atlas (HCA) portal Version 1 (Litviňuková *et al*, 2020) and Version 2 (Kanemaru *et al* 2023), which profiled the expression of 25 adult human hearts and encompasses over 700,000 cells. Relevant sample and donor information provided by the authors is reproduced in **Supplementary Table 1**. Data from this wide range of publications and donors was included to limit the potential effect of individual/technical variation and, to our knowledge, comprises the largest cross-species integrated cardiac dataset. Orthogene (Schilder and Skene, 2022) v1.8.0 was used to convert mouse gene symbols to their equivalent human orthologues via the g:Profiler database (Kolberg *et al*, 2023). Genes with ‘many-to-one’ orthologues were retained using the ‘aggregate’ function. Mouse genes without known human orthologues were excluded from analysis, but details on identity and assigned counts retained for further examination. The same approach was applied to genes from the human dataset absent from the humanised mouse data. This method retained 15,841 mouse-to-human orthologous genes, and generated human and mouse datasets directly equivalent in gene nomenclature.

### Dataset Pre-Processing and Cross-Species Integration

Celltypist (Domínguez Conde *et al*, 2022) v1.6.2, a logistic regression and machine learning-based automated celltype classifier, was used to assign cell identity using its human cardiac model (itself trained on human HCA data), and to ensure harmonious celltype nomenclature between all incorporated datasets.

Datasets were pre-processed to exclude non-expressed genes and cells with <500 total counts. Mitochondrial gene counts were excluded as they indicated a distinct technical variability between datasets. The percentage of counts mapping to ribosomal genes was calculated for each cell, and cells with >30% were excluded. Due to the extremely large size of the HCA dataset, it was downsampled to approximately 58,000 cells to match the mouse data more closely; to preserve the representation of rare celltypes, the most numerous celltypes (cardiomyocytes, fibroblasts, endothelial cells) were downsampled first, to an upper limit of 10,000 cells. Due to this downsampling procedure, and the manual pooling of isolated cells and nuclei in the preparation of samples by the original authors of respective datasets, differences in cardiac cellular proportions between species are not considered in this analysis.

Cross-species integration was then performed in R 4.2.2 (R Core Team, 2021) using Seurat 4.1.1 (Hao *et al*, 2021), with input data for CCA integration step restricted to highly-variable genes with a human-mouse orthologue expressed in both species. After integration, the standard Seurat processing and analysis workflow was applied for PCA dimension reduction, unsupervised SNN clustering and UMAP visualization. Supervised celltype annotation was then employed based on expression of canonical celltype markers across the SNN clusters to corroborate automated Celltypist annotation.

### Celltype-Specific Conserved and Differential Gene Expression

Conserved celltype marker gene detection was performed in Seurat (*FindConservedMarkers*) using the ‘roc’ method to identify and rank conserved celltype markers by specificity. The variancePartition R package (Hoffman, 2016) was used to determine the proportion of variance for each gene attributed to metadata categories of interest (celltype, species, dataset, sample).

Differential expression testing was performed in Seurat using Wilcoxon rank sum test with default parameters. Orthologous genes were tested by celltype only if expressed to a non-zero level in both species, with at least 10% of cells in either species, with differences considered statistically significant with a log2 fold-change of >= 0.58 and adjusted P-value <0.05. From this list, ‘highly divergent genes’ (HDG) were further identified as genes with a difference in percentage of cells expressing of >50% between groups. HDG lists were filtered to exclude ribosomal and mitochondrial genes.

### Downstream Analysis of Cross-Species Highly Divergent Genes

The Illuminating the Druggable Genome (IDG, Sharma and Nadler, 2021) database was queried to identify HDG classified as drug targets. Mutant Mouse Resource & Research Centers (MMRRC, Amos-Landgraf *et al*, 2022) and International Mouse Phenotyping Consortium (IMPC, Groza *et al*, 2022) databases were employed to identify HDG subject to genetic modification in mouse models. DittoSeq v1.14.0 (Bunis et al, 2020), ggplot2 (Wickham, 2016), ComplexHeatmap v2.18.0 (Gu, 2022) and scCustomize (Marsh, 2021) R packages were used for visualization.

## Results

### Mouse-to-Human Orthologue Conversion

Pre-conversion mouse data comprised 29,856 genes, which resulted in 16,112 ‘humanised’ genes after conversion to their respective human orthologue via Orthogene. These 16,112 humanised orthologues accounted for a large majority of transcript counts in the murine heart (85.8%). As mouse genes without a human orthologue were excluded from further analysis, these genes were examined to determine how their absence could affect our result; 2,718 genes had less than 3 transcript counts per heart and were not considered to contribute meaningfully to cardiac gene expression. The remaining genes tended to be of low expression, in aggregate accounting for only 14.2% of transcript counts. A considerable proportion belong to collections of relatively poorly-characterised transcripts, including pseudogenes, antisense genes and other non-coding genes predominated by ‘Gm-’ MGI (6,791) and RIKEN project (3,014) transcripts. Of the protein-coding genes in mouse which were not converted to a human orthologue, the most notable series included Olfactory receptor genes ‘Olfr^’ (361), Zinc Finger Protein ‘ZFP^’ (396), and ‘FAM’ family genes (176). Human genes in the HCA data which were absent in the humanised mouse data were also excluded from downstream analysis. These genes (16,334) accounted for 20.7% of the human transcripts, the majority of which, likewise, were non protein-coding e.g. ‘LINC-’ (2,013), or poorly characterised clone-based transcript IDs (8,987), along with gene families such as Open-Reading-Frame ‘orf’ genes (138), Zinc Finger Protein ‘ZNF^’ (323) and Immunoglobulin genes ‘IG^’ (402). Our curation of input genes facilitated a ‘like-with-like’ comparison of 15,841 human and humanised mouse genes.

### Cross-Species Integration and Cell Characterisation

After basic cell-QC filtering was applied to each dataset for minimum counts and maximum percentage of transcripts mapping to ribosomal genes, approximately 110,000 cells remained. With log-normalisation and selection of the top 3000 most variable genes, the datasets were integrated using Seurat CCA integration. Subsequently, the standard Seurat workflow was applied to the integrated assay, including PCA, UMAP and SNN clustering (**Figure 1A**). The automated annotation generated by Celltypist was supported by manual analysis of expression of canonical markers in SNN clusters. Exclusion of extremely rare celltypes with <50 cells (<0.1% of the total) in either species resulted in a cross-species integrated dataset of 16 distinct cardiac celltypes (**Figure 1B**). **Figure 1C** indicates celltype annotation by species, demonstrating a clear tendency for equivalent celltypes to co-locate in UMAP space regardless of species. As UMAP is a reduced-dimension representation of overall cellular expression, this is a *prima facie* indication of fundamental similarity in gene expression between equivalent celltypes of human and mouse heart. Yet there was a notable ‘species axis’ within some celltypes (adjacency, rather than complete overlap, of human and mouse cells of the equivalent celltype), such that sub-clusters may be largely or entirely comprised of cells from one species. By the same interpretation of UMAP this may indicate a degree of species-divergence within cognate celltypes.

**Figure 1.**
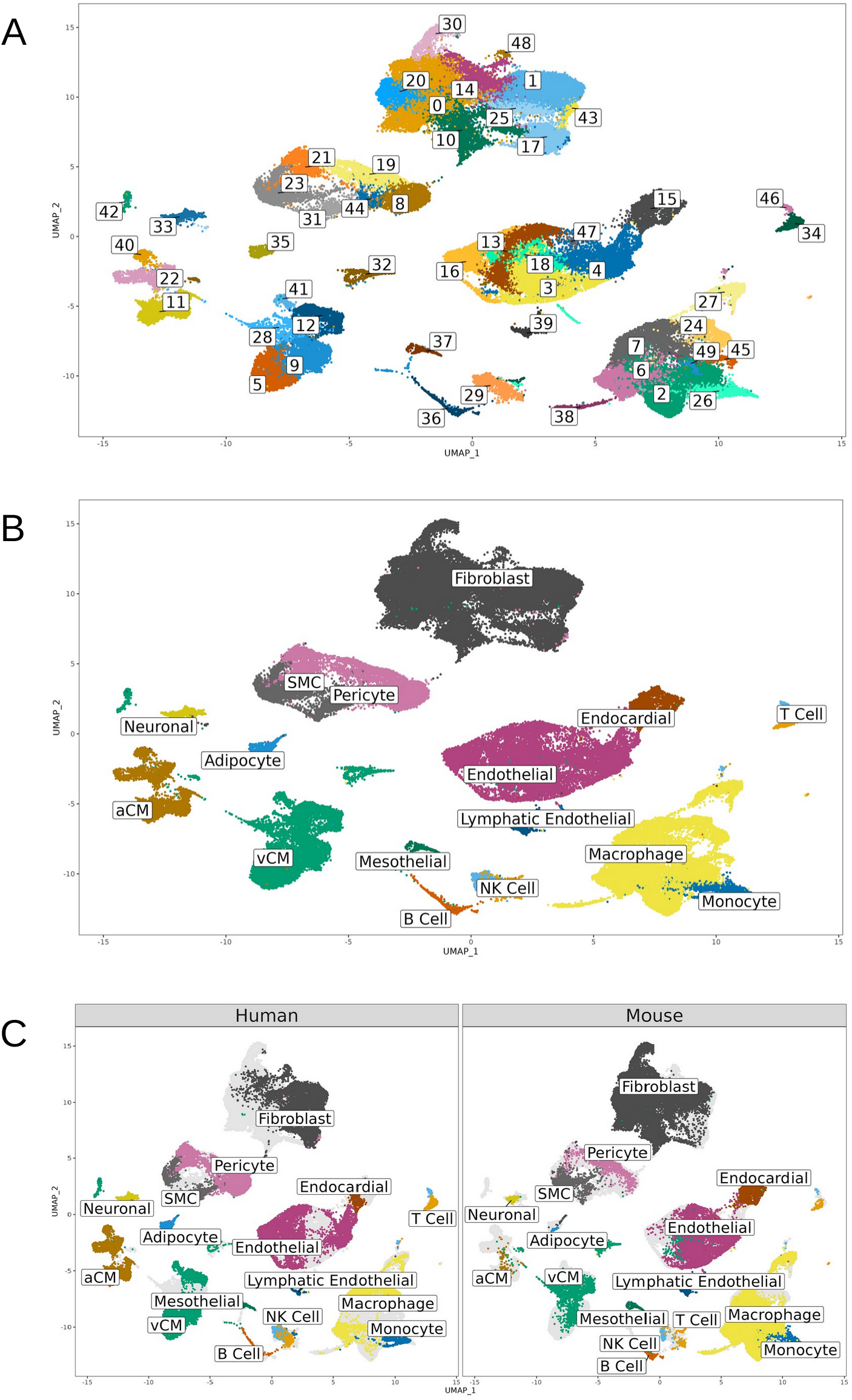
Cellular Populations in Human and Mouse Heart. mouse cardiac datasets. Distinct cellular populations are apparent, the larger of which further divide into subclusters indicating putative withincelltype heterogeneity. **B**) Combined human-mouse celltype annotation, indicating cardiac celltypes present in both species. **C**) Celltype annotation split by species of origin, confirming cells of each species contribute towards the overall celltypes.

### Transcriptomic Conservation in Human and Mouse heart

Comparative transcriptomic analyses revealed significant conservation of gene expression between all cognate celltypes of the human and mouse heart. Cells of a given type in one species were typically most similar to the equivalent celltype of the other species, and for non-immune cardiac celltypes this was always the case. For instance, mouse fibroblasts more closely resembled human fibroblasts than any other mouse celltype.

Major cardiac celltypes were identifiable in human and mouse using canonical marker genes derived from literature sources such as the Heart Cell Atlas (Litviňuková *et al*, 2020). Markers such as extracellular matrix proteins in fibroblasts (DCN, LUM, COL1A1), contractile proteins (TTN, MYH7, TNNT2) in cardiomyocytes, and adhesion proteins (PECAM1, ESAM, CDH5) in endothelial cells showed strong conservation within celltypes across species, underscoring the fundamental preservation of functional characteristics and roles of these celltypes. However, amongst some well-established celltype marker genes there were notable differences between the species (**Figure 2A**); namely COL1A1 and KCNJ8 enriched in mouse fibroblasts and pericytes respectively, and VWF, MYH7 and MYL7 enriched in human endothelial cells, vCM and aCM respectively.

**Figure 2.**
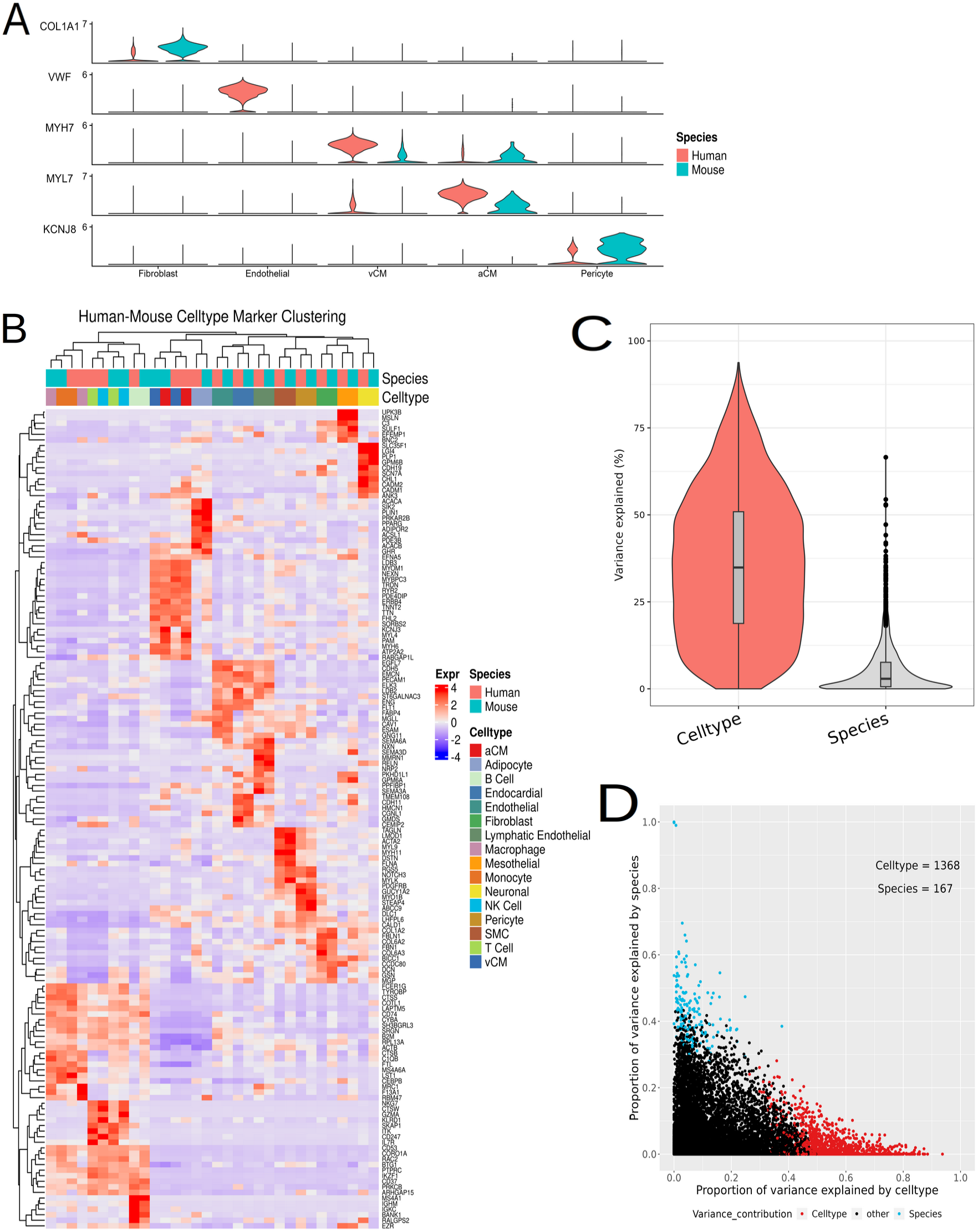
Conserved Gene Expression. **A)** Violin plot of exemplar celltype markers cardiac celltypes. Striking differences in expression between species within cognate celltypes are evident. **B**) Expression heatmap of species-conserved celltype markers – genes which distinguish a cardiac celltype across both human and mouse. **C**) Variance in gene expression across samples by Celltype and Species. Significantly greater dataset variability originates from celltype differences than species differences. **D**) Scatterplot of gene expression variance, gene (points) coloured by their largest single source of variation. Celltype-to-celltype differences were the largest factor in variance of 1368 genes, while species-to-species differences were the largest factor in only 167 genes.

Considering the potential for such inconsistency in common celltype markers, we performed an unsupervised analysis to identify cross-species celltype markers ranked by celltype classification power. We propose the use of these genes, which are agnostic to canonical celltype markers, to more reliably distinguish between celltypes in both human and mouse heart. The full list of conserved cross-species celltype markers are provided in **Supplementary Table 2**, which is intended to serve as a reference resource for celltype annotation.

Celltype-aggregated expression values of the 5 top-ranked conserved marker genes for each celltype were subjected to hierarchical clustering to assess the transcriptomic relationships between celltypes within and between species (**Figure 2B**). The main distinction occurred between immune and non-immune celltypes; immune cells tended to cluster by species rather than by celltype (i.e. human macrophage with human monocyte, mouse macrophage with mouse monocyte). Interestingly, immune cell progenitor class was recapitulated in hierarchical clusters, such that lymphoid-and myeloid-origin cells formed distinct branches. Thus, while immune celltypes tended to cluster within-species using the top-ranked conserved markers, when considered at the level of immune progenitor class our proposed markers can distinguish between celltypes across the human and mouse heart (i.e. human myeloid clusters with mouse myeloid). B cells were the notable immune cell exception, forming a distinct human-mouse cluster.

Conversely, non-immune cardiac cells exhibited a clear pattern of human-mouse clustering by celltype, such that cluster branches comprise pairs of cognate celltypes from both species (e.g. human fibroblast with mouse fibroblast). This holds true for all non-immune celltypes, demonstrating that the top conserved markers can reliably distinguish cognate celltypes between human and mouse equally. Thus, these markers reliably serve as cross-species celltype markers. A notable caveat is identified for cardiomyocyte markers; while successfully functioning as cross-species cardiomyocyte markers versus all other cell classes, the proposed markers lack sufficient specificity to distinguish between cardiomyocytes of atrial and ventricular origin by species.

In summary, our proposed cross-species celltype markers perform strongly for the major celltypes comprising cardiac tissue, but some variation between the species is noted, particularly within immune cells.

To further assess the extent of sources of variation in the dataset, gene expression was analysed with variancePartition, a statistical framework to quantify gene expression variability. For each gene, variance score was attributed to cell metadata categories such as celltype, species, sample ID and dataset ID; overall, significantly greater transcriptomic variability was attributed to celltype than species (P <2.2×10^-16^) (**Figure 2C**). Indeed, assessing each gene by the greatest single factor contributing to variance indicated 1,368 genes dominated by celltype-variability, versus only 167 for species-variability (**Figure 2C**), indicating that the transcriptomic difference between cardiac celltypes considerably outweighs the difference between human and mouse hearts. VariancePartition results are presented in **Supplementary Table 3**. Genes with variance explained predominately by celltype differences corresponded strongly with conserved celltype markers (Figure 2B, Supplementary Table 2); across all celltypes, there was an average concordance of 81.4% between species-conserved celltype markers and genes dominated by inter-celltype variation. Interestingly, this value differed between immune and non-immune cells, where average concordance of 74.12% and 84.76% was observed respectively. This appears to be aligned with the hierarchical clustering in Figure 2B, indicating relatively less celltype marker conservation among immune cells in the heart.

Additionally, genes with variance explained primarily by species differences were subsequently found to be significantly differentially-expressed between species, further explored in the following section. It follows that genes indicating both strong celltype marker conservation and low variance between species can serve as highly reliable cross-species celltype marker genes.

### Transcriptomic Divergence between Human and Mouse heart

Differential expression analysis was employed as a measure of species divergence between the celltypes of human and mouse hearts. We provide, for the first time, ranked lists of divergent gene expression between human and mouse heart, based on a direct quantitative comparison of orthologous genes within a single integrated dataset (**Supplementary Table 4**).

This analysis detected thousands of significantly differentially-expressed genes between species (**Figure 3A**), indicating transcriptomic disparity between major cardiac celltypes. While the number of differentially expressed genes is unsurprising, given the considerable underlying biological and environmental dissimilarity between the human and mouse samples, a relatively small proportion of genes exhibited extreme differences in expression. We reasoned that these genes likely contribute disproportionately to the differential responses observed between human and mouse heart. We therefore devised a method to identify a subset of ‘Highly Divergent Genes’ (HDG). These were stratified from all differentially-expressed genes based on having at least a twofold difference in expression and at least a 50 percentage-point difference in the number of cells expressing each gene within equivalent celltypes. HDGs are considered especially strong candidates of species divergence as they combine high overall expression fold-change with a considerable difference in the proportion of cell populations expressing each gene. This combination results in a clear polarization of expression between species for the 963 unique HDGs detected across all celltypes (**Figure 3B**). Visualisation of exemplar HDGs from the major cardiac celltypes (**Figure 3C-H)** confirms the remarkable contrast in expression between orthologous genes within equivalent celltypes. It is beyond the scope of this investigation to identify potential functional consequences of divergent expression in almost a thousand individual genes across 16 cardiac celltypes, and instead we focused on generating an accessible resource to facilitate other investigators employing mouse models in cross-species inquiries (**Supplementary Table 5**). Nevertheless, the HDGs are strong candidates to explain differences in function between human and mouse heart. Surprisingly, many genes prominent in cardiovascular research were listed among the HDG, and we therefore assessed how this may impact interpretation of observations in existing mouse models.

**Figure 3.**
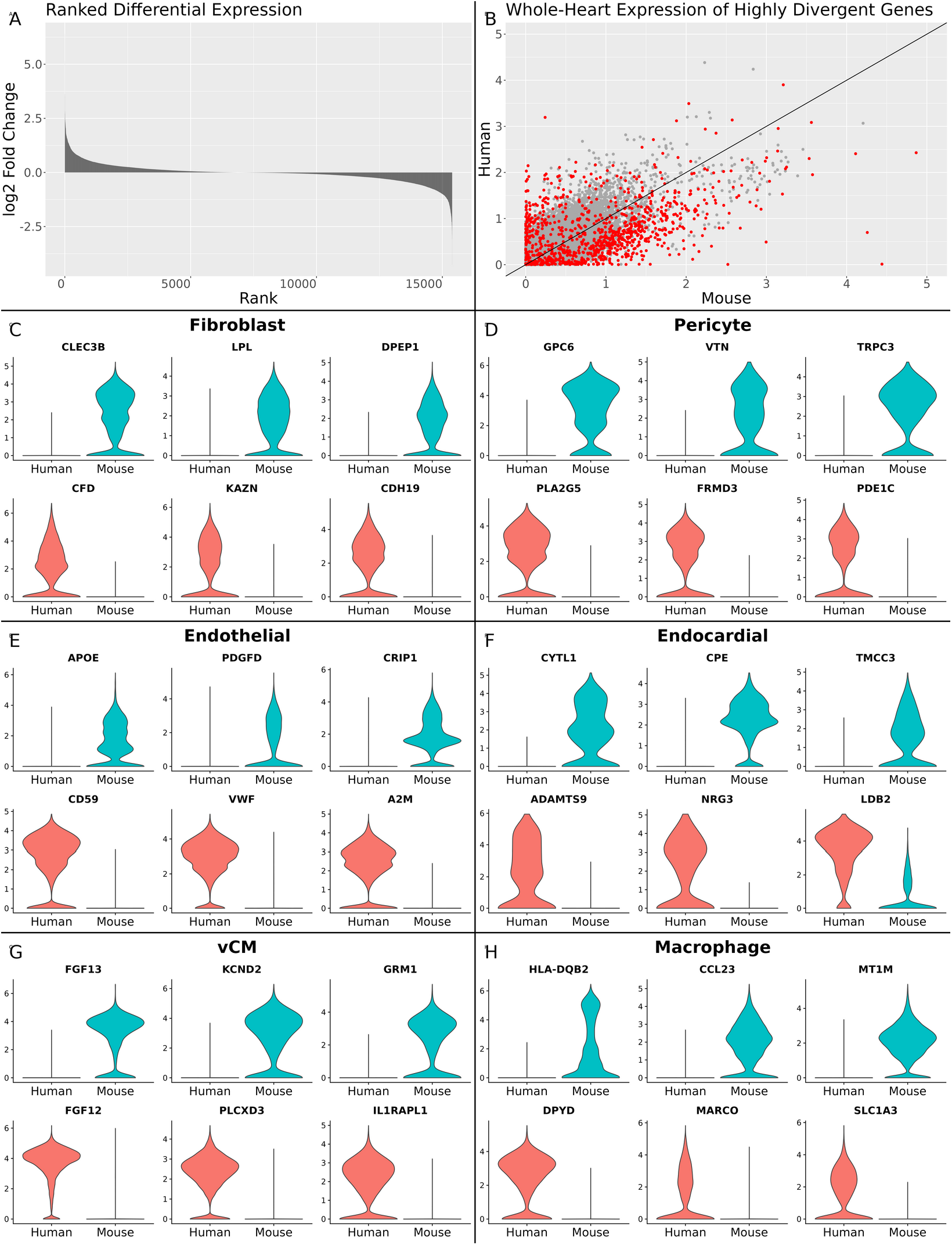
Divergent Gene Expression. **A)** Differential gene expression between human and mouse heart ranked by fold-change. Thousands of genes have expression skewed towards mouse (positive fold-change) or human (negative fold-change). **B**) Expression scatterplot of all orthologous genes, averaged across the whole human and mouse heart. Genes (points) closer to the gradient line are more conserved between species, while genes skewed towards either axis are more divergent. Points in red indicate ‘highly divergent’ genes between species in at least one cardiac celltype. **C-H**) Violin plots of highly divergent gene expression in select genes, grouped by celltype, and split by species. Such genes demonstrate extreme polarisation in expression within cognate celltypes in human and mouse heart

### Highly Divergent Genes in Gene Knockout and Drug Treatment Models

The Mutant Mouse Resource & Research Centers (MMRRC) database catalogues 24,000 unique genes with genetic modifications available in mice, of which 931 were identified as HDG between human and mouse heart. Critically, 65 of these are classified within the cardiac research category. The International Mouse Phenotyping Consortium (IMPC) catalogues gene knockouts, including 1662 with cardiovascular effects, of which 110 were HDG in our analysis (**Figure 4**).

**Figure 4.**
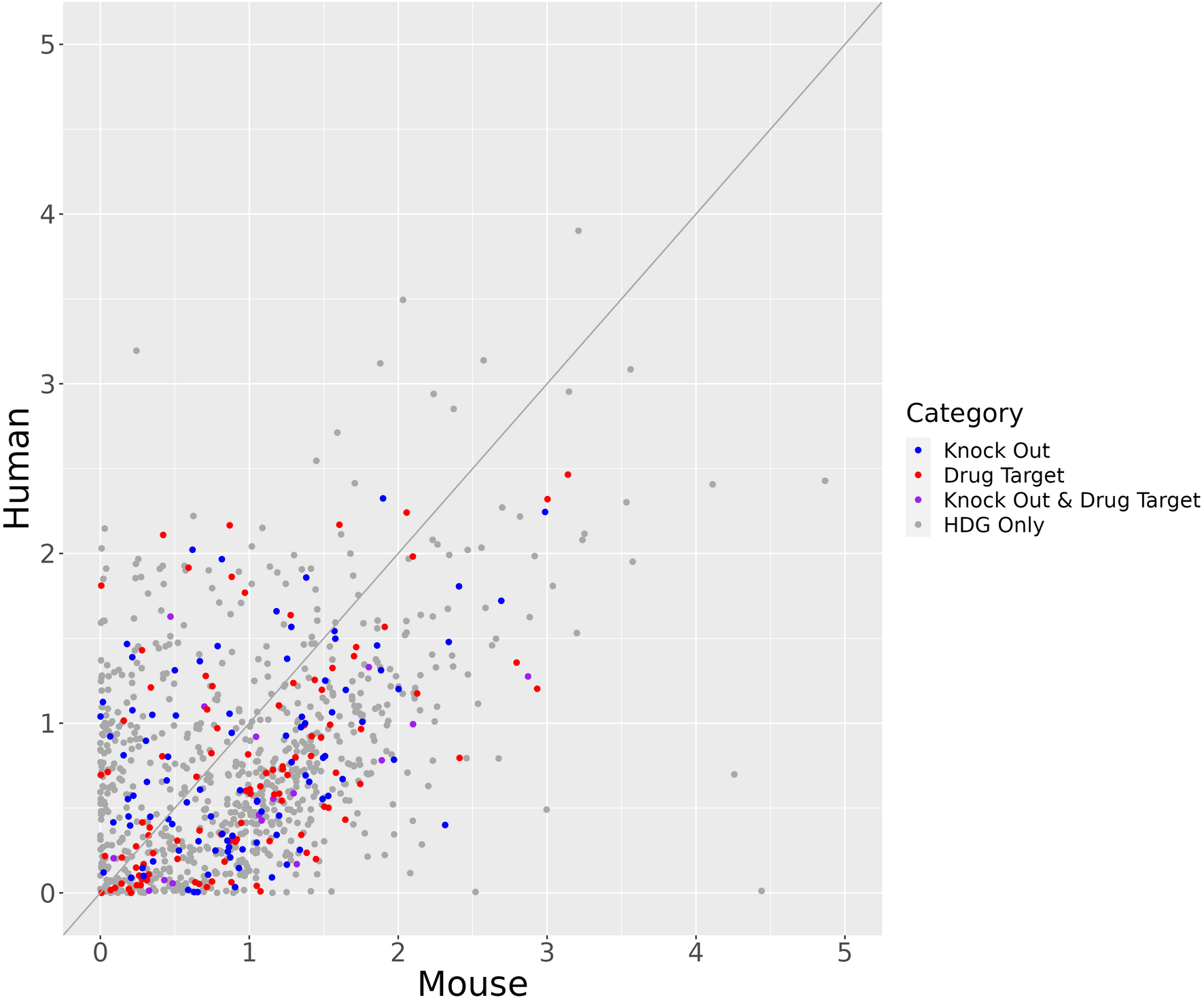
Highly Divergent Genes as Gene Knockouts and Drug Targets. Scatterplot of whole-heart expression, averaged for human and mouse whole-heart showing highly-divergent genes only. Genes (points) are coloured categorising each by knock out and drug-target status where applicable.

The Illuminating the Druggable Genome (IDG) program was queried to relate HDG with known drug target genes. 118 HDG were identified as drug targets, associated with 265 drugs in total. Drugs catalogued as targeting HDGs prominently include cardiovascular therapies such as PDE3 inhibitors (Enoximone, Amrinone, and Milrinone), Cardiac Glycosides (Acetyldigitoxin, Deslanoside, Digitoxin, Digoxin) and Angiotensin II Receptor Blocker (Losartan), with various non-cardiovascular drugs also identified including antineoplastic, antidiabetic and immunosuppressant agents.

Here we present a resource of highly-divergent genes between human and mouse heart with accompanying tags for gene knockout and drug-target information from IMPC and IDG, respectively **(Supplementary Table 6).**

## Discussion

### Human-Mouse Orthologue Conversion

In this study we directly integrated human and mouse scRNA-seq cardiac datasets, comparing the largest available resource of healthy adult human heart to five ‘humanised’ mouse heart datasets. Our cross-species integrated dataset preserved cellular expression within human-mouse orthologous genes, with the one-to-one gene orthologues accounting for a large majority of sequenced transcripts in the heart. The minority of expression data not retained in our dataset post-orthologue conversion was associated primarily with poorly-annotated or non-coding genes and transcripts. This outcome is highly similar to the findings reported by Jurado *et al* (2024) on human-mouse HF data employing similar methodology, which also focused on generating one-to-one gene orthologues to enable robust dataset integration. Thus, we determine that our orthologue-restricted dataset recapitulates biologically meaningful gene expression in both human and mouse heart, while enabling direct comparison between the constituent celltypes. Further work may be justified focused on large gene families such as olfactory receptors, zinc-finger proteins and immunoglobulins, which demonstrated rather poor human-mouse orthology and which could account for inter-species differences not assessed in our study.

### Cross-Species Celltype Proportions

While the overall celltype composition of the human and mouse heart is quite similar (Nathaly *et al*, 2022), it would be valuable to understand how compositional differences may relate to structural and, putatively, functional differences. Unfortunately, accurate assessment of *in vivo* cardiac celltype proportions was precluded in this study by the varied methodologies for tissue dissociation and sampling used by the original authors, and by the celltype downsampling method we employed, such that *in silico* cell proportions are not comparable to tissue composition. Our analysis therefore focused on gene expression, for which the data was more appropriate.

### Conservation of Celltype Marker Gene Expression

Orthologous gene expression was found to be highly conserved across human and mouse cardiac celltypes, underscoring the fundamental preservation of cellular function across mammalian tissues. Through our conserved expression analysis, we provide a cross-species marker gene resource for each cardiac celltype, accompanied by classification power in humans and mice. Ranked by celltype specificity, the top conserved genes include many ‘classical’ markers. However, it is notable that some of the most highly-ranked cross-species markers (e.g. FHL2 in cardiomyocytes, FBLN in fibroblasts, EGFL7 in endothelial cells) are much less prominent in the literature than more popular, but potentially less conserved, classical markers.

Regarding celltype-marker specificity, we observed that the top-ranked cross-species markers were particularly suited to distinguishing between non-immune cardiac cells, and less so with immune cells. This is likely due to shared expression programs within lymphoid and myeloid cell populations. Some overlap of marker genes was also evident among related classes of non-immune cells. For example, endothelial, lymphatic endothelial, and endocardial cells, in which expression of PECAM1, CDH5 and EMCN were shared, and mural cells (pericyte and smooth muscle) in which RGS5 and NOTCH3 expression was shared. This reflects physiological relatedness between classes of cells across both species, such that celltype overlap in expression of certain distinguishing genes is unsurprising. Nonetheless, this serves to emphasise the point that not all celltype markers are equally *exclusive* to a single celltype, especially when physiologically similar cells are present.

Quantification of gene expression variance between celltypes or between species, demonstrated that the overall transcriptomic difference between human and mouse heart is considerably exceeded by differences between the celltypes. Significantly greater variance was detected between individual human hearts than individual mouse hearts, despite the mouse data originating from five different research groups with a methodological, geographical and temporal spread. This serves as a reminder that underlying heterogeneity between humans is difficult to address in research using mouse strains which do not replicate the greater genetic, physiological and lifestyle diversity of humans.

### Cross-Species Transcriptomic Divergence

Perhaps most importantly, our analysis identified and ranked genes by their transcriptomic divergence in each celltype of human and mouse heart. Our analysis focused on genes for which the nature and extent of the difference means we can confidently assert their species-divergence; such genes are 1) orthologous between human and mouse, 2) expressed in heart to at least some level in both species, 3) have statistically significant differential expression, and 4) differ considerably in the percentage of each cell population expressing the gene. Such genes are extremely polarised in cross-species expression. This method identified almost one thousand genes, confidently deemed as highly species-divergent in at least one cardiac celltype, a resource which to our knowledge has not previously been reported.

Although we do not assess in detail the functional consequences of these differences, it is possible that physiological divergence between human and mouse heart is related to the findings we describe here. For instance, cases of near-total polarisation of genes in the same celltype were detected – in such cases, contrasting protein levels and subsequent pathway functions may be expected, unless an equivalent but non-orthologous gene can fulfil the role of the ‘missing’ gene.

The obvious application of this resource is in planning of cross-species cardiac research. A preliminary search for conservation or divergence in genes or celltypes under investigation would require trivial effort, while offering considerable benefit (i.e., ruling out lines of investigation unlikely to be validated across species, enabling focus on more promising candidates). Retrospective analysis of existing data is also viable to enhance understanding of failure to replicate findings across species.

### Interpretation of HDG in Genetically Modified Mouse Models

The fact that almost all HDGs reported here have been subject to genetic modification in mouse, including at least 110 with gene knockouts causing reported cardiovascular effects, is intriguing. Knocking out (or otherwise modifying) a highly-expressed gene in mouse may be of limited value in modelling human biology if the same gene is normally absent or expressed at extremely low levels in the equivalent human cells. Although the *phenotype* resulting from genetic modification is typically the primary aim of such models, our work indicates that greater care should be taken to understand the cross-species celltype-specific nature of gene expression, and the potential ramifications of over-interpreting physiological/phenotype changes in human-mouse cardiac research resulting from genetic modifications of highly-divergent genes. This issue is of potentially even greater concern regarding genetic modification in cell lines; while divergent gene expression in a celltype *in vivo* could putatively be compensated for by expression of the same gene in another celltype, this is not possible in cell monoculture. Thus, where human cell lines are intended to validate changes in mouse models or *vice versa*, particular care should be taken to ascertain that the intended genetic modification will target a gene normally expressed at appreciable levels in that celltype in both species.

### Interpretation of HDG in Drug Target Genes

A similar issue is raised concerning HDGs encoding drug targets. This is a complex topic, considering; 1) levels of gene expression and gene product follow only a general correlation which can vary considerably between genes and celltypes (Kosti *et al*, 2016), 2) gene products may be secreted and transported to different cells than those expressing the gene, or to extracellular regions, and 3) drug targets may have localised or systemic distribution. Each of these factors may differ between human and mouse. Therefore, we do not claim to predict cross-species drug efficacy using this resource. Nonetheless, where the expression of a known drug-target gene is found to have considerable cross-species difference, there are possible downstream ramifications on drug activity. The drug targeting information we include for our species-divergent genes may therefore prove useful in research on comparative pharmacological effects of therapies in human versus mouse.

### Conclusion

Direct integration of scRNA-seq datasets from human and mouse hearts using orthologous genes facilitated a comprehensive comparative assessment of the transcriptomes of cognate cardiac celltypes from each species. The high degree of transcriptomic conservation observed among cardiac celltypes highlights the fundamental similarity in celltype gene expression between humans and mice. Nonetheless, we uncovered nearly one thousand genes exhibiting clear divergence in expression between species.

The observation that many of the highly divergent genes have been targeted in mouse models or considered as drug targets underscores the importance of understanding cross-species differences in gene expression for translational research. Moreover, the observed heterogeneity among human individuals serves as a reminder of the challenges in extrapolating findings from mouse models to human populations.

Single-cell transcriptomics has highlighted the importance of considering celltype-specific differences in expression of orthologous genes between species. This information has important implications for the design and interpretation of cross-species cardiac research studies.

## Acknowledgements

We are grateful for the public availability of gene expression data from human and mouse heart.

